# Transcriptional Downregulation of a Type III Secretion System under Reducing Conditions in *Bordetella pertussis*

**DOI:** 10.1101/2020.04.20.052324

**Authors:** Masataka Goto, Tomoko Hanawa, Akio Abe, Asaomi Kuwae

## Abstract

*Bordetella pertussis* uses a type III secretion system (T3SS) to inject virulence proteins into host cells. Although the *B. pertussis* T3SS was presumed to be involved in host colonization, the efficient secretion of type III secreted proteins from *B. pertussis* has not been observed. To investigate the roles of type III secreted proteins during infection, we attempted to optimize culture conditions for the production and secretion of a type III secreted protein, BteA, in *B. pertussis*. We observed that *B. pertussis* efficiently secretes BteA in ascorbic acid-depleted (AsA^−^) medium. When L2 cells, a rat lung epithelial cell line, were infected with *B. pertussis* cultured in the AsA^−^ medium, BteA-dependent cytotoxicity was observed. We also performed an immunofluorescence assay of L2 cells infected with *B. pertussis*. The clear fluorescence signals of Bsp22, a needle structure of T3SS, were detected on the bacterial surface of *B. pertussis* cultured in the AsA^−^ medium. Since ascorbic acid is known as a reducing agent, we cultured *B. pertussis* in liquid medium containing other reducing agents such as 2-mercaptoethanol and dithioerythritol. Under these reducing conditions, the production of type III secreted proteins was repressed. These results suggest that in *B. pertussis*, the production and secretion of type III secreted proteins are downregulated under reducing conditions.

**IMPORTANCE:** The type III secretion system (T3SS) of *Bordetella pertussis* forms a needle-like structure that protrudes from the bacterial cell surface. *B. pertussis* uses T3SS to translocate virulence proteins called effectors into host cells. The culture conditions for effector production in *B. pertussis* have not been investigated. We attempted to optimize culture medium compositions for producing and secreting type III secreted proteins. We found that *B. pertussis* secretes type III secreted proteins in reducing agent-deprived liquid medium, and that BteA-secreting *B. pertussis* provokes cytotoxicity against cultured mammalian cells. These results suggest that redox signaling is involved in the regulation of *B. pertussis* T3SS.

## Introduction

The genus *Bordetella* is made up of Gram-negative bacteria and has been subdivided into 16 subspecies to date (1). *Bordetella pertussis* is a causative agent of whooping cough, also known as pertussis. *B. pertussis* strictly adapts to humans, and it lacks an environmental reservoir (2). A variety of virulence factors have been identified in *Bordetella*, e.g., filamentous hemagglutinin (3), adenyl cyclase toxin (4) and pertactin (5). These virulence factors are regulated by a *Bordetella* two-component regulatory system, BvgAS. In addition to these virulence factors, the type III secretion system (T3SS) of *B. pertussis* is also regulated by BvgAS (6, 7).

A T3SS that forms a needle-like structure is conserved among many Gram-negative bacteria. In order to subvert hosts, bacteria use a T3SS to inject type III secreted proteins into host cells (8). Such type III secreted proteins are called effectors. Three types of effectors have been identified in *Bordetella*: BteA ((9) [also referred to as BopC (10)], BopN (11, 12), and BspR (13) [also referred to as BtrA (14, 15)]. BteA is 658 amino acids in length and is localized to lipid rafts in host cells via its N-terminal region (16). BteA induces necrotic cell death in several types of cultured mammalian cells (9, 10), and BteA translocation into host cells is promoted by the functioning of BopN (11). BspR is a negative regulator for type III secreted proteins (13) and is translocated into nuclei of cultured mammalian cells (17). BspR has been suggested to be an anti-sigma factor against BtrS that functions as a positive regulator for type III secreted proteins (15). Some type III secreted proteins are called translocators rather than effectors. Three types of translocators have been identified in *Bordetella*, i.e., BopB (18), BopD (19) and Bsp22 (20), and they function as the path for effectors delivered into host cells. The complex of BopB and BopD forms the pores on the host membrane (19), and Bsp22 forms a filamentous-like structure and is associated with the T3SS needle and with a pore-forming component, BopD (21).

The optimal culture conditions for the production and secretion of BteA have not been established. To investigate the roles of BteA and other type III secreted proteins during *B. pertussis* infection, we investigated culture conditions for the maximal production and secretion of type III secreted proteins in *B. pertussis* strain Tohama I.

## MATERIALS AND METHODS

### Bacterial strains

The strains used in this study are listed in Table 1. *B. pertussis* Tohama I (22) was used as the wild-type strain. BP344, BP348, BP350, and BP351 were used as clinical isolates and were a kind gift from Dr. K. Kamachi (National Institute of Infectious Diseases) (23). *Escherichia coli* DH10B and Sm10λ*pir* were used for DNA cloning and conjugation, respectively (Table 1). *B. pertussis* was grown on Bordet-Gengou agar plates at 37°C for 5 days. Bordet-Gengou agar plates were prepared as follows. We mixed 15 g of Bordet-Gengou agar base (BD #248200), 5 mL of glycerol, 100 mL of defibrinated horse blood (Nihon Bio-Supp Center), and 400 mL of distilled water. The mixture was autoclaved and poured onto plastic plates. Fresh colonies of *B. pertussis* on each Bordet-Gengou agar plate were suspended in Stainer-Scholt (SS) medium [Table 2, (24)]. In this study, we used a liquid medium described as a cyclodextrin-containing medium as the SS medium (25). Low casamino acids (LCA) medium was prepared by removing the agar from cyclodextrin solid medium (CSM) contents [Table 2, (26)]. *B. pertussis* was cultured in SS medium with a starting A_600_ of 0.23 under static conditions at 37°C. To measure the bacterial density, a spectrophotometer (Ultrospec 2100 *pro*, Amersham) was used. The final A_600_ of *B. pertussis* Tohama I culture after 22 hr was ca 0.67, and the culture contained ca 1.0 x 10^9^ cfu/mL. The liquid cultivation period was 22 hr for the protein preparation, mRNA preparation, and infection.

**Table 1.** Bacterial strains and plasmids used in study

| Strains or plasmids | Relevant genotype/description | Reference |
| --- | --- | --- |
| Strains |  |  |
| *B. pertussis* |  |  |
| Tohama I | Clinical isolate | 22 |
| BP344 | Clinical isolate | 23 |
| BP348 | Clinical isolate | 23 |
| BP350 | Clinical isolate | 23 |
| BP351 | Clinical isolate | 23 |
| MGKU1 | Tohama I nalidixic-acid resistant strain | This study |
| MGKU2 | T3SS-inactive strain | This study |
| MGKU3 | BteA-deficient strain | This study |
| MGKU4 | BspR-deficient strain | This study |
| *E. coli* |  |  |
| DH10B | Host strain for pDONR201 | invitrogen |
| Sm10λpir | Host strain for pABB-CRS2 | 18 |
| Plasmids |  |  |
| pDONR201 | DNA cloning vector, Km<sup>r</sup> | invitrogen |
| pABB-CRS2 | Suicide vector for conjugation, Amp<sup>r</sup> | 27 |
| pABB-$\Delta bscN$ | pABB-CRS2, $\Delta bscN$ gene | 18 |
| pABB-$\Delta bopC$ | pABB-CRS2, $\Delta bopC$ ($\Delta bteA$) gene | 10 |
| pMGKU401 | pDONR201, *bspR* gene | this study |
| pMGKU402 | pDONR201, $\Delta bspR$ gene | this study |
| pMGKU403 | pABB-CRS2, $\Delta bspR$ gene | this study |

**Table 2.** Medium compositions

| Basal medium (g/L) | SS | LCA |
| --- | --- | --- |
| Sodium Gultamate | 10.7 | 10.7 |
| L-proline | 0.24 | 0.24 |
| NaCl | 2.5 | 2.5 |
| KCl | 0.2 | 0.2 |
| KH<sub>2</sub>PO<sub>4</sub> | 0.5 | 0.5 |
| MgCl<sub>2</sub> •6H<sub>2</sub>O | 0.1 | 0.1 |
| CaCl<sub>2</sub> | 0.02 | 0.02 |
| Tris | 6.1 | 6.1 |
| Casamino acids | 10 | 0.5 |
| Dimethyl-β-cyclodextrin | 1 | 1 |

| Supplement (g/L) | SS | LCA |
| --- | --- | --- |
| L-cysteine•HCl | 4 | 4 |
| FeSO<sub>4</sub>•7H<sub>2</sub>O | 1 | 1 |
| L-ascorbic acid | 40 | 2 |
| Niacin | 0.4 | 0.4 |
| L-glutathione reduced | 15 | 15 |

### Generation of a Tohama I nalidixic acid-resistant mutant

For the generation of a Tohama I nalidixic acid-resistant mutant, Tohama I cultured in SS medium for 24 hr as described above was spread on 20 sets of Bordet-Gengou agar plates containing nalidixic acid at the final concentration of 30 *μ*g/ml. After 5 days, surviving colonies were designated as MGKU1. The growth rate, colony size and hemolysis on the BG plates of the resulting nalidixic acid-resistant strain were almost equivalent to those of the wild-type strain, and we observed that the amounts of the production and secretion of type III secreted proteins in MGKU1 were not significantly different from those of Tohama I wild-type (data not shown).

### Cell culture

L2 cells, a rat lung epithelial cell line (ACTT CCL-149), were maintained in F-12K (Invitrogen, Tokyo). The cell culture medium was supplemented with 10% fetal calf serum (FCS). L2 cells were grown at 37°C under a 5% CO_2_ atmosphere.

### Construction of gene-disrupted mutants

The primers used in this study are listed in Table 3. To construct the *bspR* mutant, the 4.0 kb DNA fragment encoding *bspR* and its flanking regions was amplified by polymerase chain reaction (PCR) with primers B1-bspR and B2-bspR using *B. pertussis* Tohama I genomic DNA as the template. The resulting PCR product was cloned into pDONR201 to obtain pMGKU401, by means of adapter PCR and site-specific recombination techniques using a Gateway cloning system (Invitrogen). An inverse PCR was carried out with the set of primers R1-bspR and R2-bspR using circular pMGKU401 as the template.

**Table 3.**
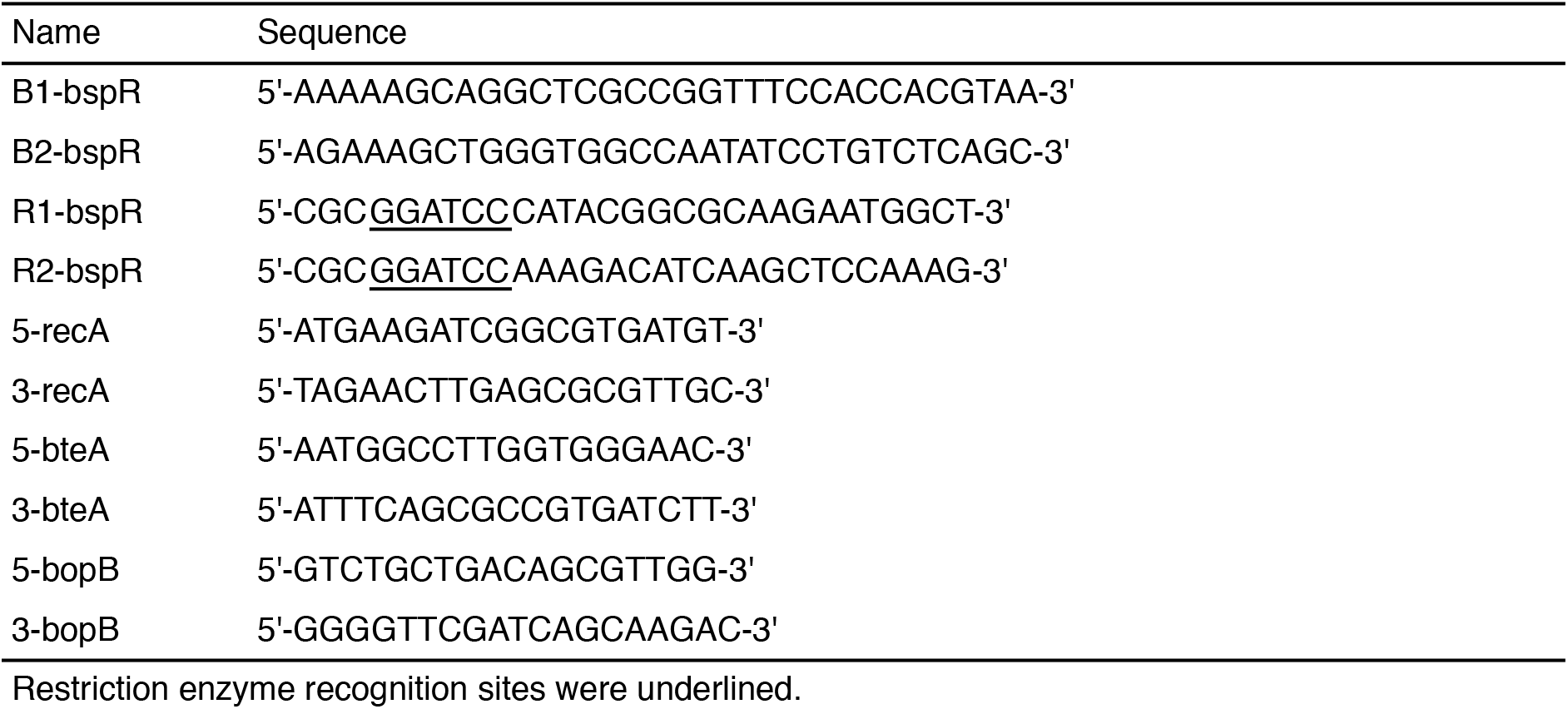
Primers used in this study.

The resulting PCR product was digested with BamHI and self-ligated to obtain pMGKU402. This plasmid contained the BamHI site in addition to a 474-bp in-frame deletion from 175 bp downstream of the 5’ end of the *bspR* gene to 52 bp upstream of the 3’ end of the gene. This plasmid, pMGKU402, was mixed with a positive suicide vector, pABB-CRS2 (27), to obtain pMGKU403 using the Gateway cloning system. This plasmid, pMGKU403, was introduced into *E. coli* Sm10λ*pir* (18) and transconjugated into MGKU1 as described previously (28). The resulting mutant strain was designated as MGKU4 (BspR-deficient strain). Similarly, *pABB*-Δ*bscN* (18) and pABB-*ΔbopC* (10) were separately introduced into *E. coli* Sm10λ*pir* and were transconjugated into MGKU1. The resulting mutant strains were designated as MGKU2 (a T3SS-inactive strain) and MGKU3 (a BteA-deficient strain), respectively.

### Preparation of proteins from culture supernatants and whole bacterial cell lysates

Secreted proteins released into bacterial culture supernatants and whole bacterial cell lysates were prepared by trichloroacetic acid precipitation. In the case of the supernatant fraction (Sup) samples, the culture supernatants were filtered with a 0.22 *μ*m filter (Millipore #SLGVR33RB). Next, 2 *μ*L of 5% deoxycholic acid and 100 *μ*L of 100% trichloroacetic acid were added to the 1 mL of filtered supernatants. After being incubated on ice for 15 min, the samples were centrifuged at 21130 ×*g* for 5 min. The resulting precipitated proteins were neutralized with 2 *μ*L of 2 M Tris-base and dissolved in 13 *μ*L of 2×SDS-PAGE sample buffer. In the case of the whole cell lysate (WCL) samples, bacterial pellets collected from 1 mL of bacterial cultures were resuspended in distilled water. 100 *μ*L of 100% trichloroacetic acid was added to 1 mL of the bacterial suspension. After being incubated on ice for 15 min, the samples were centrifuged at 21130 ×*g* for 5 min. The resulting precipitated proteins were neutralized with 5 *μ*L of 2 M Tris-base and dissolved in 95 *μ*L of 2×SDS-PAGE sample buffer. When the A_600_ of bacterial culture was 0.67, 0.5 *μ*L of WCL and 5 *μ*L of Sup samples were loaded on the SDS-PAGE gels. The loaded sample amounts were adjusted by the A_600_ of each bacterial culture in order to load samples prepared from the same number of bacteria. The protein samples were separated by SDS-PAGE and analyzed by western blotting.

### Antibodies

Anti-BteA, anti-BopB, and anti-Bsp22 antibodies were purified from rabbit serum in our previous study (10, 18, 29). To detect filamentous hemagglutinin (FHA) signals, we used mouse anti-FHA serum (30). Mouse anti-CyaA monoclonal antibody was purchased from Santa Cruz Biotechnology (Santa Cruz, CA). To prepare the anti-BtrS antibody, the peptide corresponding to the C-terminus region of BtrS (CALREALRERGYDSVP) were conjugated with hemocyanin from keyhole limpets (Sigma) by using 3-maleimidobenzoic acid N-hydroxysuccinimide ester (Sigma). These cross-linked peptides were used to immunize rabbits, and resulting anti-sera were incubated with peptide immobilized on epoxy-activated sepharose 6B (Amersham) to obtain specific Ig-fractions.

### Quantitative reverse transcription-PCR (qRT-PCR)

The amounts of mRNA were measured by qRT-PCR. Bacterial total RNA was prepared using a Trizol Max Bacterial RNA Isolation Kit (Invitrogen), an RNeasy Mini Kit (Qiagen, Hilden, Germany) and an RNase-free DNase-free Kit (Qiagen). The RNA sample was reverse transcribed with Transcriptor Universal cDNA Master (Roche Diagnostics, Indianapolis, IN) and a T100 Thermal Cycler (Bio-Rad, Hercules, CA). The resulting cDNA was amplified by FastStart Essential DNA Probes Master (Roche) using the following primer pairs: 5-recA and 3-recA for *recA*; 5-bteA and 3-bteA for *bteA*; 5-bopB and 3-bopB for *bopB* (Table 3). The results were calculated as described in the Roche manual. The amount of *recA* mRNA was used as an internal control. The mRNA amounts are presented herein as relative to the amounts in the Tohama I cultured in SS medium, which was defined as 1.

### LDH Assays

To examine the release of lactate dehydrogenase (LDH) from *B. pertussis-infected* cells, 5.0×10^4^ cells/well of L2 cells seeded in 24-well plates were infected with bacteria at the multiplicity of infection (MOI) of 125, 250, or 500. The plates were centrifuged at 900 *g* for 5 min and incubated for 3 hr at 37°C under a 5% CO_2_ atmosphere. The amounts of LDH were measured spectrophotometrically using a Cyto-Tox 96 Non-radioactive Cytotoxicity Assay Kit (Promega, Madison, WI). We added 50 *μ*L of 10 %Triton X-100 to the 950 *μ*L of extracellular medium of L2 cells in 24-well plate. We then mixed the extracellular medium by pipetting. We subtracted the LDH value obtained from extracellular medium of uninfected cells from the value obtained from the Triton X-100-treated cells and used the resulting value as 100%.

### Immunofluorescent staining

For the immunofluorescent staining assay, 2.5×10^5^ cells/well of L2 cells were seeded on coverslips in six-well plates and incubated overnight, and then infected with *B. pertussis* at an MOI of 125. The plates were centrifuged at 900 *g* for 5 min and incubated for 3 hr at 37°C under a 5% CO_2_ atmosphere. The infected L2 cells were then immunostained as described previously (12). Briefly, the cells were treated with 4% paraformaldehyde, 50 mM NH4Cl, 0.2% Triton-X, and 4% bovine serum albumin (BSA). After blocking, the cells were stained by anti-Bsp22 antibody (29). As a secondary antibody, Alexa Fluor 488 goat anti-rabbit IgG (Invitrogen) was used. F-actin was stained with Rhodamine Phalloidin (Molecular Probes, Eugene, OR). Bsp22 signals per one cell were counted under a fluorescence microscope.

### Statistical analyses

The statistical analyses were performed using the nonparametric unpaired *t*-test with a one-tailed p*-*value with Prism ver. 5.0f software (Graphpad, La Jolla, CA). Values of p<0.05 were considered significant.

## RESULTS

### Type III secreted proteins are secreted from *B. pertussis* cultured in LCA medium

Cyclodextrin solid medium (CSM) is reported as a synthetic medium for clinical isolation of *B. pertussis* (26). We sought to determine whether *B. pertussis* grown on this agar secretes type III effectors, and we prepared a liquid medium, designated “low casamino acids (LCA)” medium, by removing the agar from CSM contents (Table 2). *B. pertussis* was cultured in SS or LCA medium, and then the whole cell lysates (WCL) and supernatant fractions (Sup) samples were subjected to western blotting with antibodies against BteA (an effector, a type III secreted protein) and BopB (a translocator, a type III secreted protein).

BteA and BopB signals were detected in WCL and Sup of the wild-type Tohama I cultured in LCA medium (Fig. 1). As reported previously, BteA forms SDS-resistant multimers, and in our present study we also detected the signals at around 200 kDa. In contrast, the BteA and BopB signals were faintly detected in the WCL samples of SS medium, and no BteA and BopB signals were detected in the Sup samples of MGKU2 (a T3SS-inactive strain) or the wild-type cultured in SS medium (Fig. 1). These results suggest that *B. pertussis* secretes type III secreted proteins in LCA medium.

**Fig. 1.**
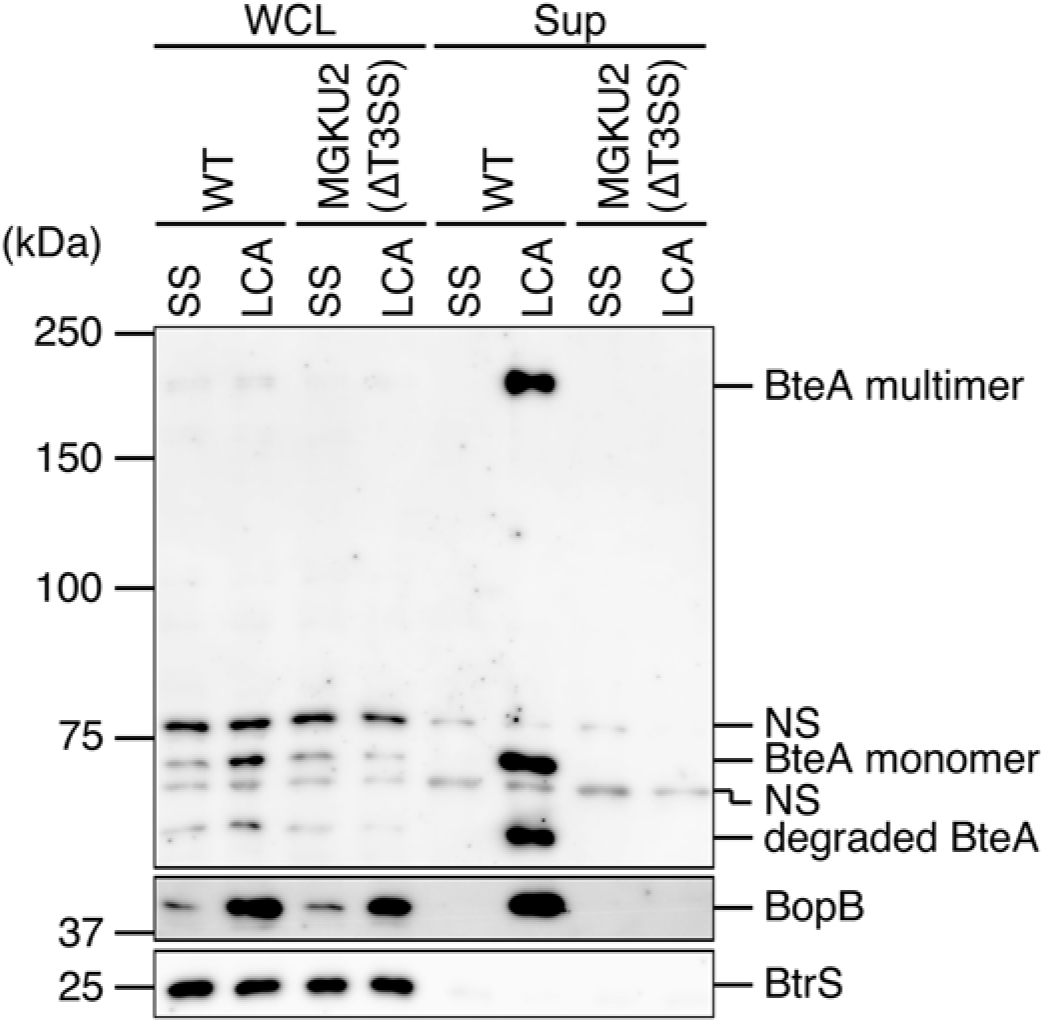
BteA and BopB production by *B. pertussis* Tohama I cultured in LCA medium. Tohama I and MGKU2 (a T3SS-inactive strain) were cultured in SS or LCA medium. Whole cell lysates (WCL) and supernatant fraction (Sup) samples were separated by SDS-PAGE and analyzed by western blotting with antibodies against BteA (*upper panel*), BopB (*middle panel*) or BtrS (*lower panel*). BtrS, an RNA polymerase sigma factor (15), was used as the unsecreted protein control. BteA forms SDS-resistant multimers (10). NS: nonspecific signals. Loaded WCL and Sup samples were prepared from an equal volume of bacterial culture. Experiments were performed at least three times, and representative data are shown.

### Ascorbic acid downregulates the production and secretion of type III secreted proteins in *B. pertussis*

Although the ingredients of the LCA and SS media are the same, the amounts of ascorbic acid and casamino acids in LCA medium are lower than those in SS medium (Table 2). To determine which substance is involved in the production and secretion of type III secreted protein, we cultured *B. pertussis* in SS medium, ascorbic acid-deprived SS medium (SS_AsA^−^), casamino acid-deprived SS medium (SS_CaA^−^), or both ascorbic acid- and casamino acid-deprived medium (SS_AsA^−^_CaA^−^). WCL and Sup samples were subjected to western blotting with antibodies against BteA and BopB (Fig. 2).

**Fig. 2.**
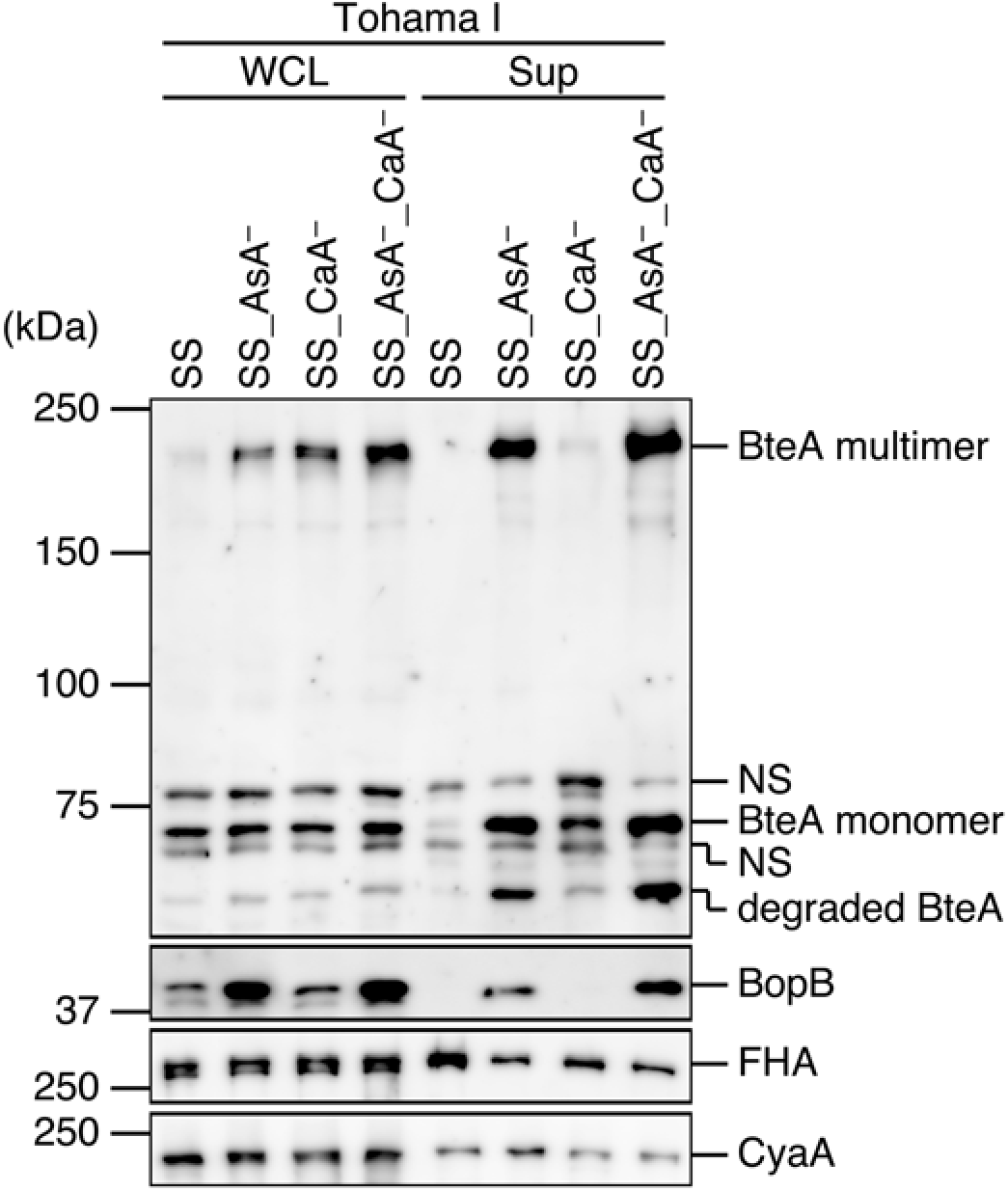
Effects of ascorbic acid and casamino acids on Bvg-regulated virulence factors. Tohama I was cultured in SS medium or ascorbic acid-deprived SS medium (SS_AsA^−^), casamino acid-deprived SS medium (SS_CaA^−^) or both ascorbic acid- and casamino acid-deprived SS medium (SS AsA^−^_CaA^−^). WCL and Sup samples were separated by SDS-PAGE and analyzed by western blotting with antibodies against BteA, BopB, FHA, or CyaA. NS: nonspecific signals. Loaded WCL and Sup samples were prepared from an equal volume of bacterial culture. Experiments were performed at least three times, and representative data are shown.

The BteA multimer signal intensities in the WCL samples of the SS_AsA^−^, SS_CaA^−^ and SS_AsA^−^_CaA^−^ media were stronger than that in the SS medium (Fig. 2). A faint BteA signal intensity in the Sup samples of SS was detected (Fig. 2). The BteA signal intensities in the Sup samples were evident in the SS_AsA^−^ and SS_AsA^−^_CaA^−^ media (Fig. 2). Again, BopB signal intensities in both WCL and Sup samples were evident in the SS_AsA^−^ and SS_AsA^−^_CaA^−^ media (Fig. 2).

We also investigated the potential involvement of ascorbic acid or casamino acids in Bvg-regulated virulence factors, e.g., FHA and CyaA. WCL and Sup samples were subjected to western blotting with antibodies against FHA and CyaA (Fig. 2). The results revealed that there were no significant differences in the FHA or CyaA signal intensities in the WCL and Sup samples among the media used here (Fig. 2). Collectively, these results suggest that ascorbic acid has a specific influence on the production and secretion of BteA and BopB in *B. pertussis*.

Standard SS medium contains ascorbic acid at a final concentration of 2270 *μ*M. To further explore whether ascorbic acid affects the production and secretion of type III secreted proteins, we cultured *B. pertussis* in SS medium in the presence of ascorbic acid at the final concentrations of 91, 454, or 2270 *μ*M. WCL and Sup samples were prepared from each bacterial culture and analyzed by western blotting (Fig. 3). The BteA signal was detected in both WCL and Sup samples of SS medium at 0–454 *μ*M ascorbic acid, but the BteA signal of the SS medium at 2270 *μ*M ascorbic acid was faint or absent (Fig. 3). The BopB signals showed a similar pattern (Fig. 3). These results indicate that *B. pertussis* produces and secretes type III secreted proteins under low ascorbic acid concentrations.

**Fig. 3.**
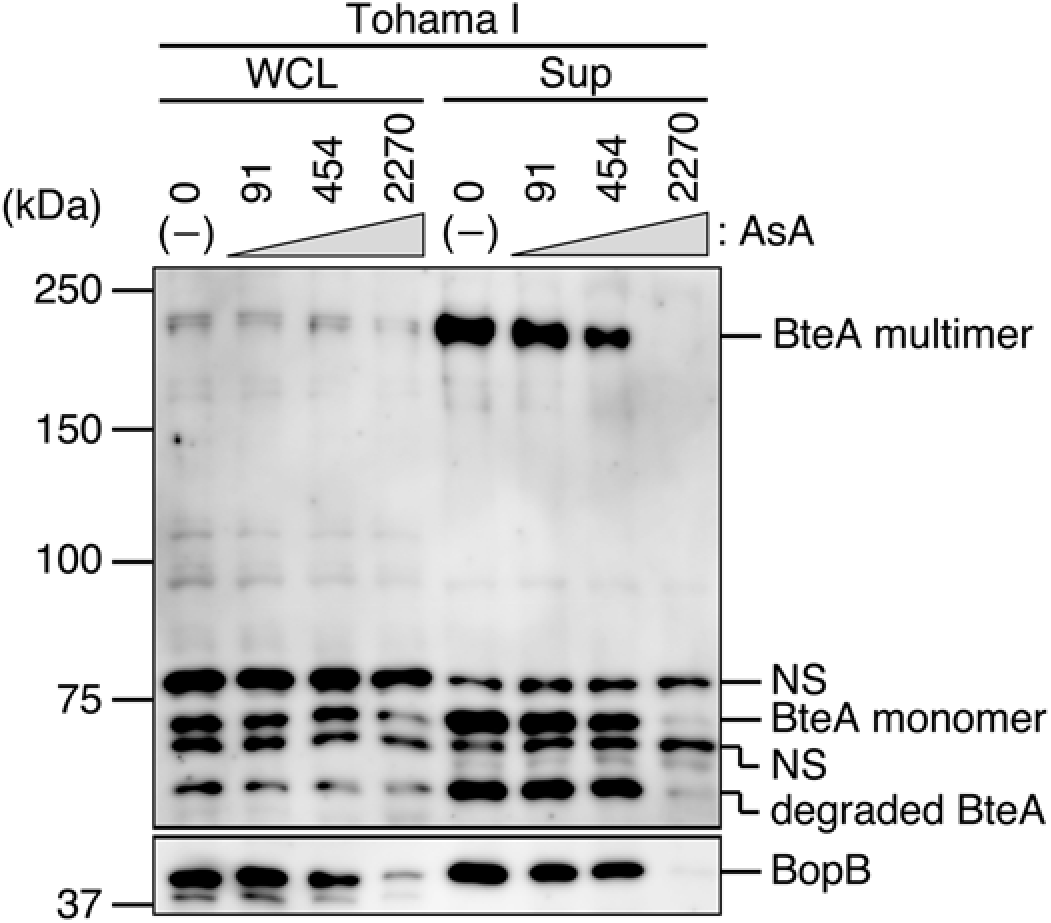
Effects of ascorbic acid on type III secreted proteins. Whole cell lysates and secreted proteins prepared from Tohama I cultured in SS medium in the presence of ascorbic acid at the final concentration of 91, 454 or 2270 *μ*M were separated by SDS-PAGE and then analyzed by western blotting with antibodies against BteA or BopB. NS: nonspecific signals. Loaded WCL and Sup samples were prepared from an equal volume of bacterial culture. Experiments were performed at least three times, and representative data are shown.

### BteA and BopB are secreted from other *B. pertussis* clinical isolates cultured in SS_AsA^−^ medium

To determine whether *B. pertussis* clinical isolates secrete type III secreted proteins under low ascorbic acid concentrations, we cultured four strains of *B. pertussis* (Table 1) in SS medium or ascorbic acid-deprived medium (SS_AsA^−^). The prepared WCL and Sup samples were analyzed by western blotting (Fig. 4). BteA signals in the Sup samples of BP350 and BP351 were evident in the SS_AsA^−^ medium (Fig. 4, AsA^−^). Again, BopB signals in the Sup samples of BP350 and BP351 were evident in SS_AsA^−^ medium. These results suggest that some clinical strains, e.g., BP350 and BP351, also secrete type III secreted proteins under low ascorbic acid concentrations.

**Fig. 4.**
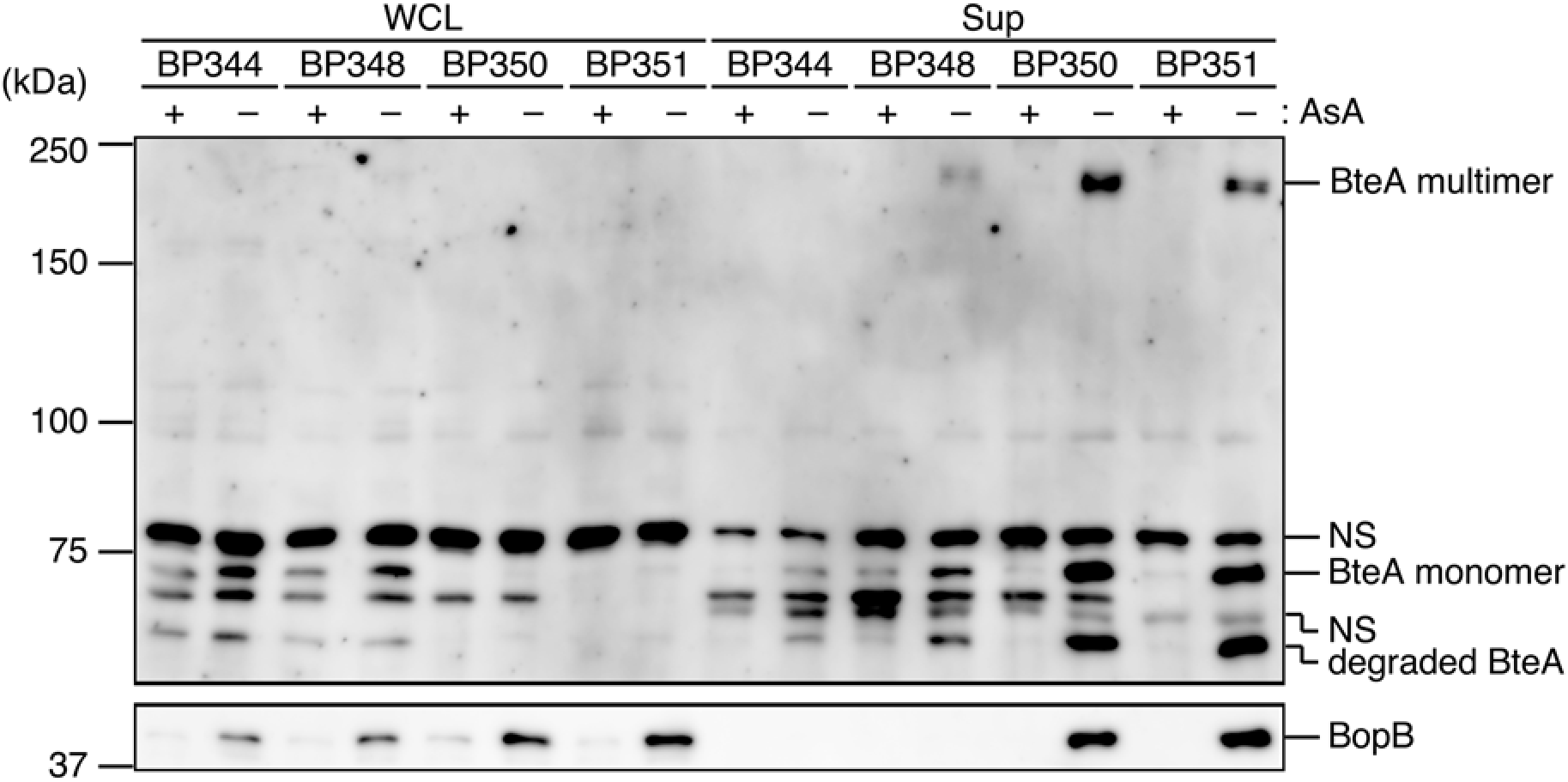
Effects of ascorbic acid on the secretion of BteA and BopB in clinical isolates cultured in SS_AsA^−^ medium. The *B. pertussis* clinical isolates BP344, BP348, BP350, and BP351 were cultured in SS (+) or SS_AsA^−^ (−) medium. WCL and Sup samples were separated by SDS-PAGE and analyzed by western blotting with antibodies against BteA (*upper panel*) or BopB (*lower panel*). NS: nonspecific signals. Loaded WCL and Sup samples were prepared from an equal volume of bacterial culture. Experiments were performed at least three times, and representative data are shown.

### Translocator BopB is upregulated at the transcriptional level in *B. pertussis* cultured in SS_AsA^−^ medium

To investigate the gene expressions of type III secreted proteins under ascorbic acid-starved conditions, we prepared total RNA from *B. pertussis* cultured in SS or SS_AsA^−^ medium. The cDNA samples reverse-transcribed from the total RNA samples were subjected to a quantitative RT-PCR analysis to quantify the relative amounts of *bteA* and *bopB* mRNA. The results demonstrated that the relative amount of *bteA* mRNA of Tohama I in SS_AsA^−^ medium was increased compared to that in SS medium (Fig. 5A). However, the relative amount of *bteA* mRNA of BP350 in the SS_AsA^−^ medium was not significantly different from that in the SS medium (Fig. 5A). The relative amounts of *bopB* mRNA of Tohama I and BP350 in SS_AsA^−^ medium were increased compared to those in SS medium (Fig. 5B).

**Fig. 5.**
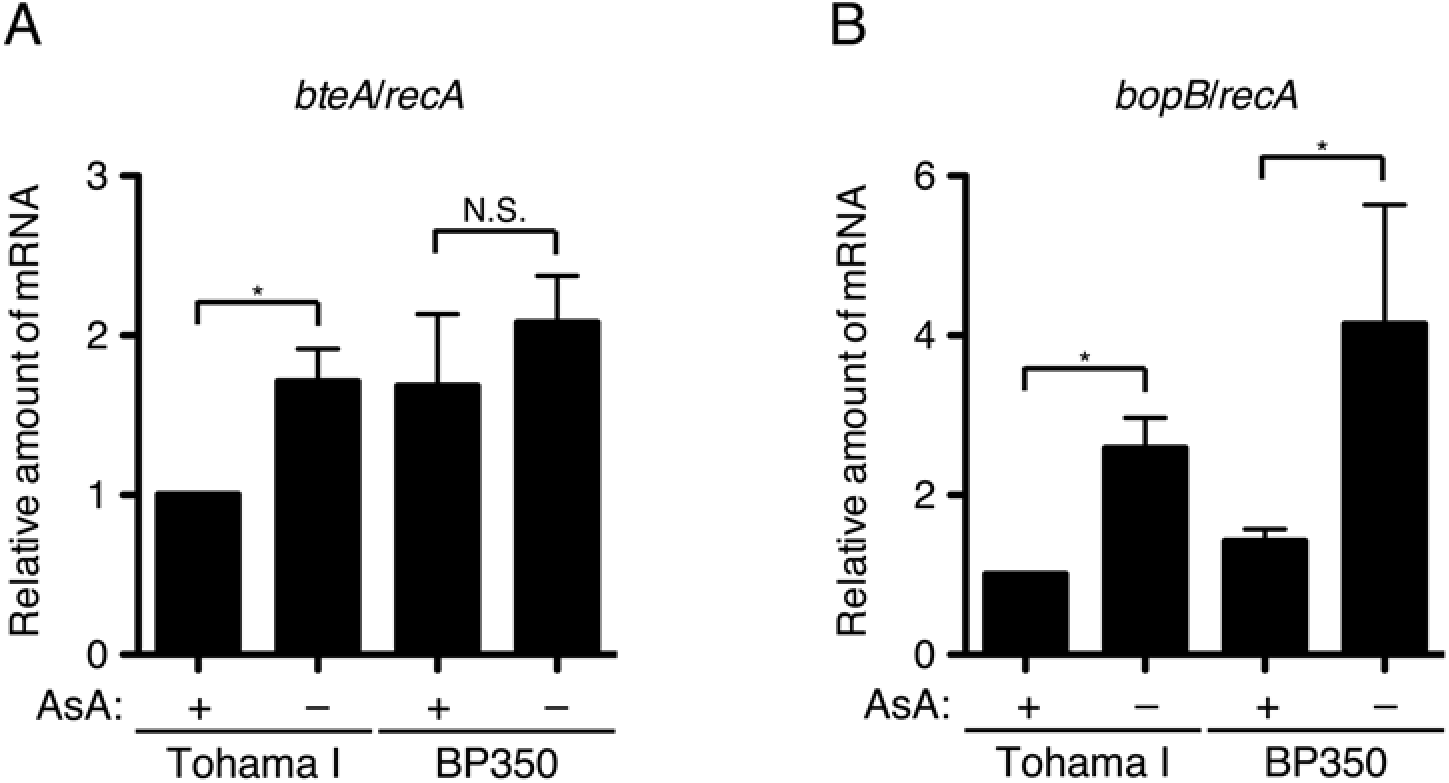
Results of the RT-PCR analysis for mRNA levels of *bteA* and *bopB* from *B. pertussis* cultured under ascorbic acid-starved conditions. Total RNA was prepared from Tohama I or BP350 cultured in SS or SS_AsA^−^ medium and subjected to a quantitative RT-PCR analysis. The histograms show the relative amounts of *bteA* (**A**) and *bopB* (**B**) mRNA normalized by the housekeeping gene, *recA* mRNA. Error bars: the standard error of the mean (SEM) from triplicate experiments. *p<0.05; N.S.: no significant difference. Experiments were performed at least three times, and representative data are shown.

These results suggest that BopB is upregulated at the transcriptional level in Tohama I and BP350 under low ascorbic acid concentrations. BteA is also upregulated at the transcriptional level in Tohama I under low ascorbic acid concentrations but not in BP350.

### *B. pertussis* induces cytotoxicity against cultured mammalian cells

A *B. pertussis* BspR-deficient strain was reported to induce cytotoxicity against cultured mammalian cells (15). In the present study, we identified better culture conditions for the production and secretion of BteA in *B. pertussis* than those provided with standard SS medium. We thus investigated whether wild-type *B. pertussis* provokes cytotoxicity against cultured mammalian cells under these conditions. L2 cells, a rat lung epithelial cell line, were infected at an MOI of 500 for 3 hr in several modified SS media shown in Fig. 6. The amounts of ascorbic acid and/or casamino acids was reduced to one-twentieth of and/or to one-half of those in SS medium. We also used the SS media without ascorbic acid and/or casamino acids.

**Fig. 6.**
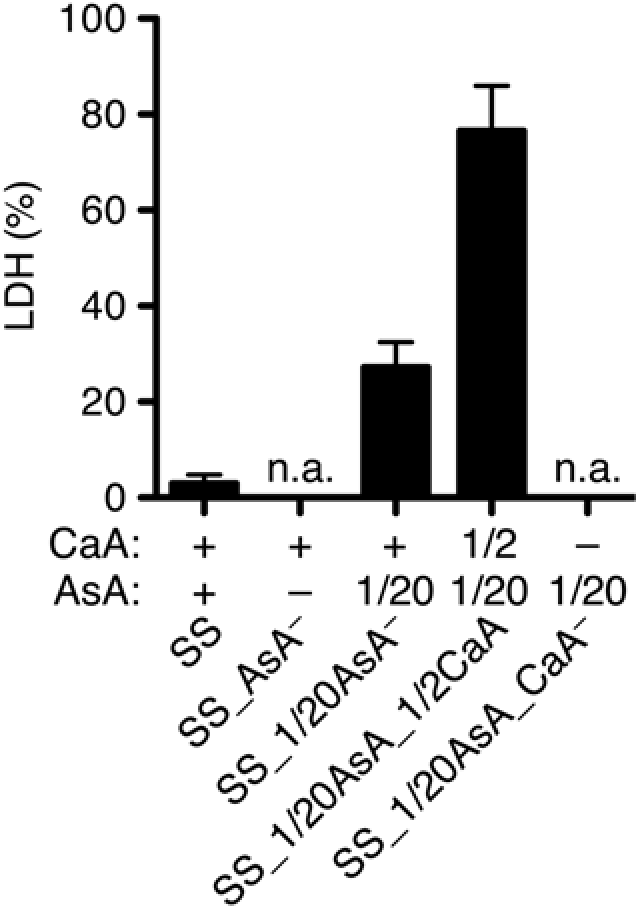
Cytotoxic activity of *B. pertussis* cultured in SS_AsA^−^ medium. L2 cells were infected with Tohama I cultured in SS medium, SS_AsA^−^ medium, SS medium containing one-twentieth of ascorbic acid (SS_1/20AsA), SS medium containing one-twentieth of ascorbic acid and one-half of casamino acids (SS_1/20AsA_1/2CaA), or SS medium containing one-twentieth of ascorbic acid and no casamino acids (SS_1/20AsA_CaA^−^) at an MOI of 500 for 3 hr. +, 1/2 and 1/20 show the amounts of indicated substances that are equal to, one-half of, and one-twentieth of those in SS medium, respectively. The amounts of LDH released into the extracellular medium from infected cells are shown, and the relative cytotoxicity (%) was determined as described in the Materials and Methods section. Error bars: the SEM from triplicate experiments. n.a.: No available result. Experiments were performed at least three times, and representative data are shown.

The amounts of lactate dehydrogenase (LDH) released into the extracellular medium were then measured.

The results revealed that released LDH was detected from L2 cells infected with Tohama I in the SS_1/20AsA or SS_1/20AsA_1/2CaA medium (Fig. 6). The SS_1/20AsA medium and SS_1/20AsA_1/2CaA medium showed no cytotoxicity. On the other hand, the L2 cells incubated in the SS_AsA^−^ or SS_1/20AsA_CaA^−^ medium were injured by the medium itself. Therefore, these media were not useful for this assay. Taken together, these results demonstrate that we successfully observed cytotoxicity of mammalian cells induced by wild-type *B. pertussis*.

### The T3SS of *B. pertussis* cultured in SS_AsA^−^ medium is active during infection against cultured mammalian cells

As described above, we observed cytotoxicity of *B. pertussis* against cultured cells.

In order to confirm that *B. pertussis* indeed secretes type III secreted proteins during infection, we infected L2 cells with *B. pertussis* in SS or SS_1/20AsA*_*1/2CaA medium at an MOI of 125 for 3 hr. After infection, Bsp22 (a translocator, and a component of the filamentous-like structure) and F-actin were stained with anti-Bsp22 and rhodamine phalloidin, respectively.

The results showed that the amounts of Bsp22 signals (Fig. 7A, green) on L2 cells infected with Tohama I and BP350 cultured in SS_1/20AsA_1/2CaA medium were greater than those in SS medium (Fig. 7). Thus, we demonstrated that *B. pertussis* T3SS is activated during the infection of cultured mammalian cells under appropriate conditions.

**Fig. 7.**
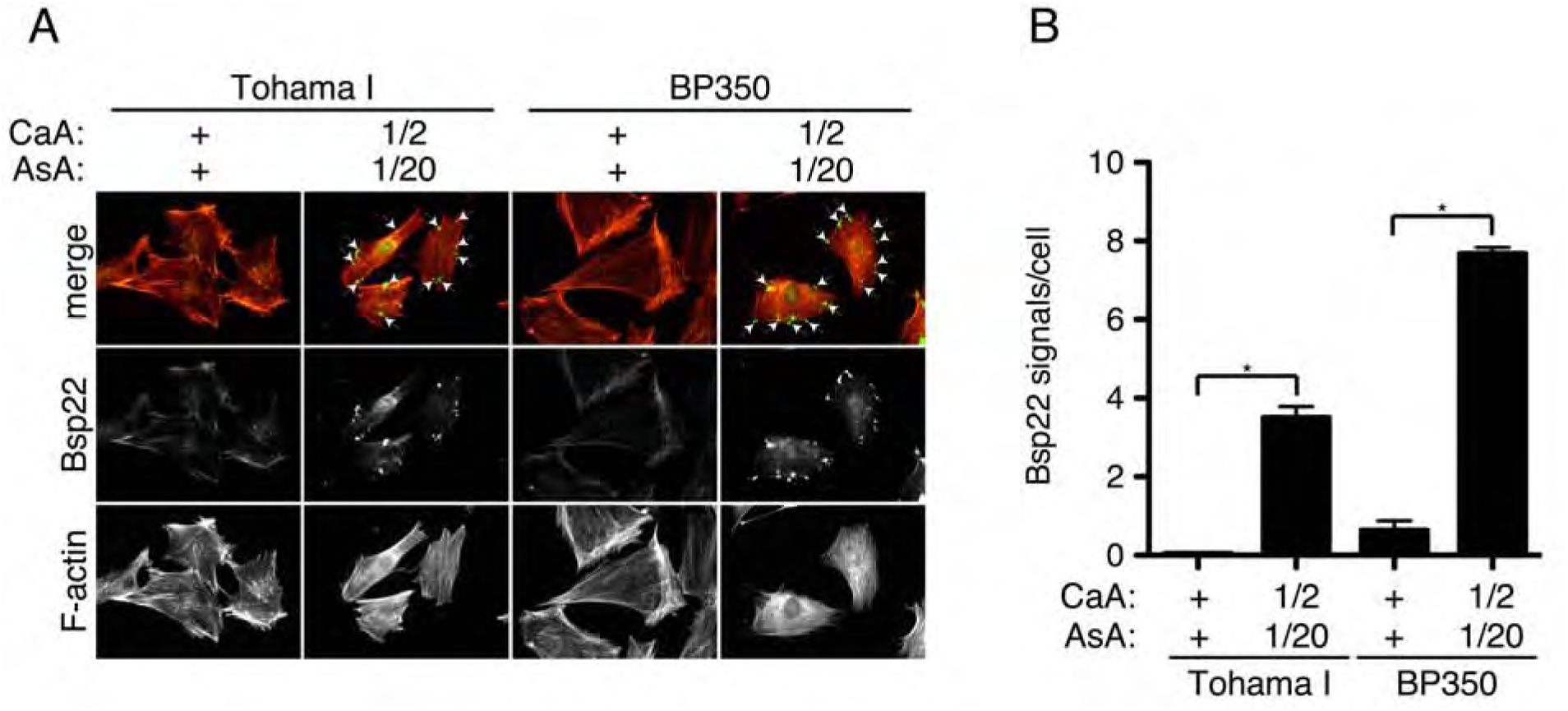
Immunofluorescent staining of Bsp22 on L2 cells infected with *B. pertussis*. **A:** L2 cells were infected with Tohama I or BP350 in the indicated media at an MOI of 125 for 3 hr. +, 1/2 and 1/20 indicate the amounts of indicated substances that are equal to, one-half of, and one-twentieth of those in SS medium, respectively. After fixation, cells were stained with anti-Bsp22 antibody (*green*) and rhodamine phalloidin (*red). Arrowheads* indicate the Bsp22 (*green*) signal. **B:** Bsp22 signals per one cell were counted under a fluorescent microscope. At least 120 cells were randomly chosen. Error bars: the SEM from triplicate experiments. *p<0.05; N.S.: no significant difference. Experiments were performed at least three times, and representative data are shown.

### The cytotoxicity induced by wild-type *B. pertussis* depends on the T3SS

It has been reported that *B. bronchiseptica* induces BteA-dependent necrotic cell death against cultured mammalian cells (10). To investigate whether the cytotoxicity provoked by wild-type *B. pertussis* also depends on BteA function, we infected L2 cells with MGKU1 (a Tohama I nalidixic acid-resistant strain), MGKU2 (a T3SS-inactive strain), MGKU3 (a BteA-deficient strain), or MGKU4 (a BspR-deficient strain that allows the constitutive activation of T3SS) in SS_1/20AsA_1/2CaA medium at an MOI of 125 or 250 for 3 hr. The amounts of LDH released into extracellular medium were measured.

As a result, LDH was not detected in the medium of L2 cells infected with MGKU2 (Fig. 8A,B). The amount of LDH detected in the medium of L2 cells infected with MGKU3 was significantly lower than that in the medium of L2 cells infected with MGKU1 (Fig. 8A,B). The amount of LDH detected in the medium of L2 cells infected with MGKU4 was significantly greater than that in the medium of L2 cells infected with MGKU1 at an MOI of 125 (Fig. 8A), but not at an MOI of 250 (Fig. 8B).

**Fig. 8.**
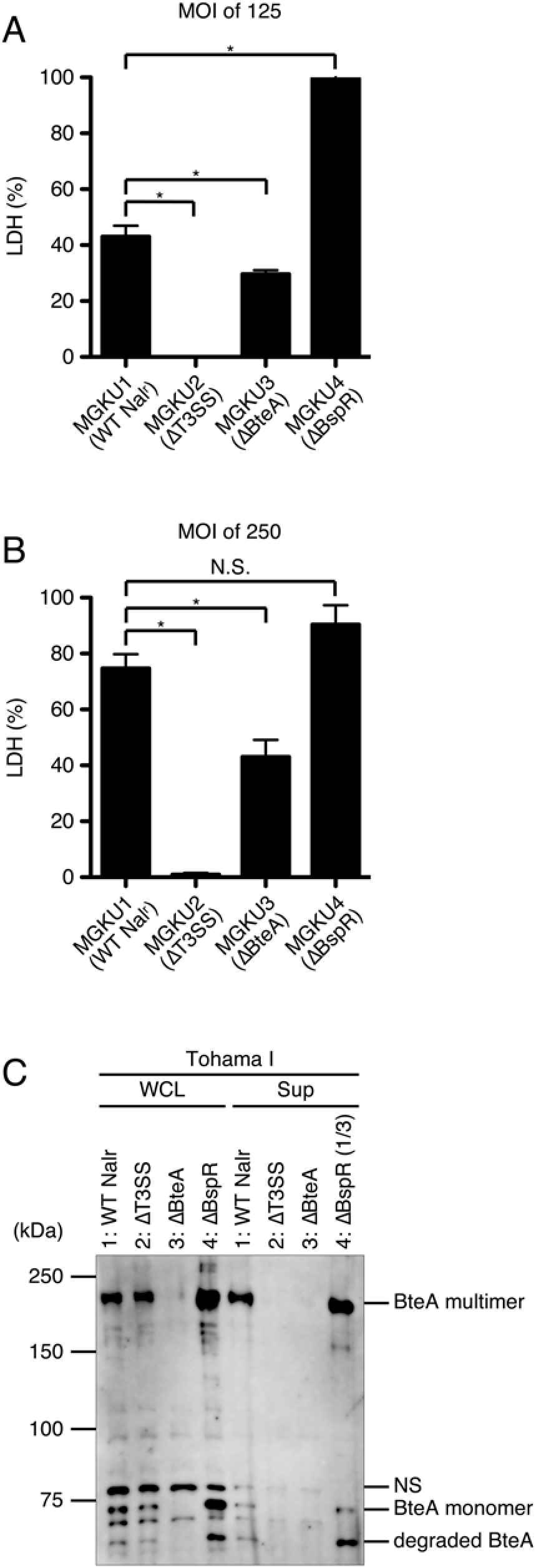
Cytotoxic activities of *B. pertussis* mutants. The results of LDH assays at an MOI of 125 (**A**) or 250 (**B**) are shown as a histogram. Error bars: the SEM from triplicate experiments. *p<0.05. Experiments were performed at least three times, and representative data are shown. **C:** MGKU1 (Tohama I nalidixic acid-resistant strain, 1: WT Nal^r^), MGKU2 (T3SS-inactive strain, 2: ΔT3SS), MGKU3 (BteA-deficient strain, 3: ΔBteA), or MGKU4 (BspR-deficient strain, 4: ΔBspR) was cultured in SS_1/20AsA_1/2CaA medium. WCL and Sup samples were separated by SDS-PAGE and analyzed by western blotting with anti-BteA antibody. We loaded a 3-fold smaller amount of MGKU4 supernatant fraction on the SDS-PAGE gel to avoid obtaining an excess signal intensity. NS: nonspecific signals. Loaded WCL and Sup samples were prepared from an equal volume of bacterial culture. Experiments were performed at least three times, and representative data are shown.

WCL and Sup samples were prepared from each strain and separated by SDS-PAGE, then analyzed by western blotting with anti-BteA antibody (Fig. 8C). BteA signals of the WCL samples were detected in MGKU1, MGKU2, and MGKU4, but not in MGKU3 (Fig. 8C). BteA signals of the Sup samples were detected in MGKU1 and MGKU4, but not in MGKU2 or MGKU3 (Fig. 8C). These results suggest that the cytotoxicity induced by wild-type *B. pertussis* (Fig. 6) is dependent on T3SS activity and is partially dependent on BteA.

### The secretion of type III secreted proteins in *B. pertussis* is downregulated under reducing conditions

Ascorbic acid is known as a reducing agent. To determine whether the secretion of type III secreted proteins is downregulated under reducing conditions, we cultured Tohama I in ascorbic acid-deprived SS medium (SS_AsA^−^, Fig. 9, “none”) and in SS_AsA^−^ medium containing the reducing agents ascorbic acid (AsA), 2-mercaptoethanol (2-ME) or dithioerythritol (DTE), respectively. The prepared WCL and Sup samples were analyzed by western blotting with anti-BteA, anti-BopB, or anti-FHA antibodies (Fig. 9). The BteA and BopB signal intensities of both WCL and Sup samples were decreased in the presence of each of the reducing agents (Fig. 9). In contrast, the FHA signal intensities of both WCL and Sup samples were not affected by the reducing agents. The growth of bacteria cultured in SS medium with reducing agents was not significantly different from that of the bacteria cultured in SS medium (data not shown). These results suggest that the production and secretion of type III secreted proteins in *B. pertussis* are downregulated under reducing conditions.

**Fig. 9.**
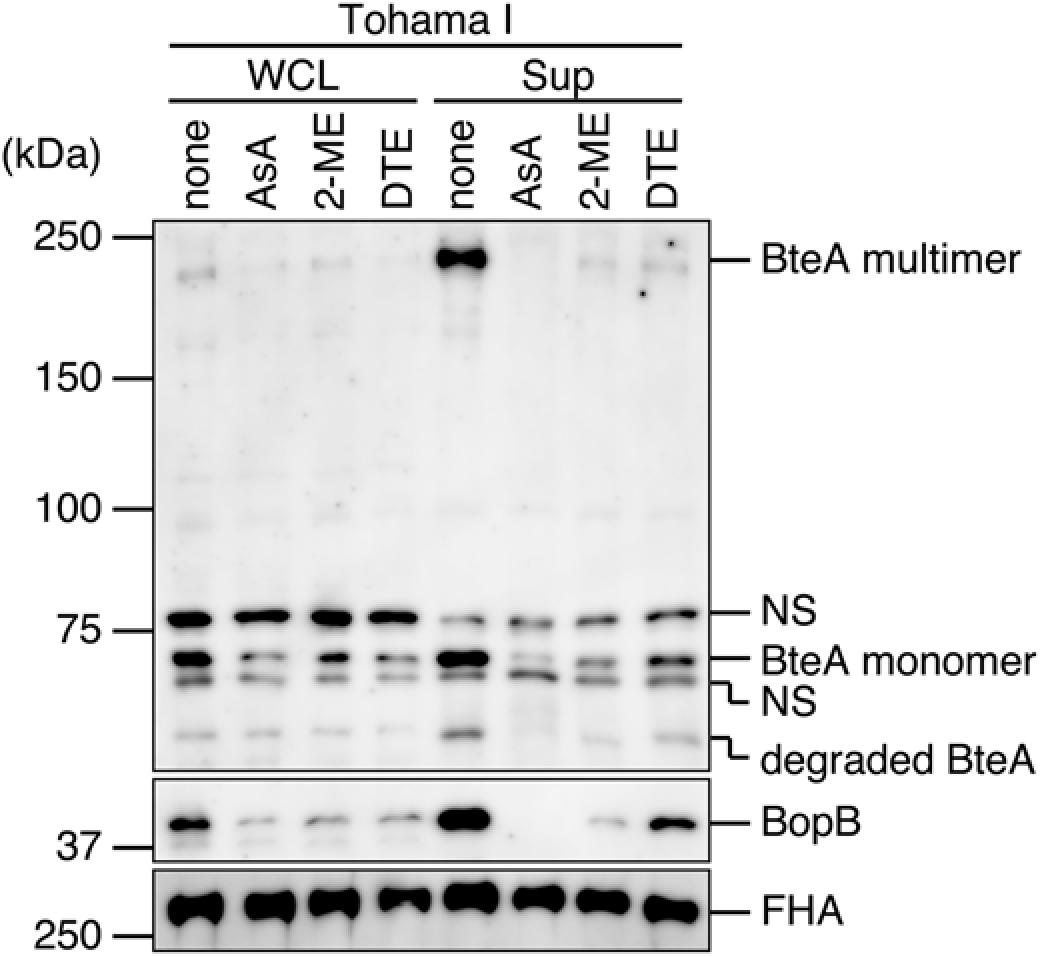
Effects of reducing agents on the production and secretion of type III secreted proteins in *B. pertussis*. Tohama I was cultured in SS_AsA^−^ medium or SS_AsA^−^ medium containing ascorbic acid (AsA), 2-mercaptoethanol (2-ME), or dithioerythritol (DTE) at final concentrations of 2.3, 2.1, or 3.0 mM, respectively. WCL and Sup samples were separated by SDS-PAGE, then analyzed by western blotting with antibodies against BteA, BopB, or FHA. Loaded WCL and Sup samples were prepared from an equal volume of bacterial culture. NS: nonspecific signals. Experiments were performed at least three times, and representative data are shown.

## DISCUSSION

In this study, *B. pertussis* increased the production and secretion of type III secreted proteins, BteA and BopB, when cultured in ascorbic acid-deprived SS medium (SS_AsA^−^). We successfully produced cytotoxicity against mammalian cells by the infection of wild-type Tohama I cultured in SS 1/20AsA_1/2CaA medium.

*B. pertussis* decreased the production and secretion of type III secreted proteins when cultured in SS_AsA^−^ medium containing reducing agents such as 2-ME or DTT. These results suggest that redox signaling is involved in regulation of the T3SS in *B. pertussis*.

It has been reported that no secretion of the type III secreted proteins BteA and Bsp22 was detected from Tohama I (31), and that the secretion of the type III secreted proteins BteA and BopD was detected from a non-vaccine-type strain, BP157. However, BP157 showed no cytotoxicity against L2 cells, J774 mouse macrophage-like cells, and HeLa cells (32). In another report, glutamate limitation upregulates production of the type III secreted proteins (33). However, the requirement of a type III secretion system (T3SS) for virulence in *B. pertussis* thus remains unclear. On the other hand, a BscN-deficient mutant in which the T3SS was inactive colonized mouse lungs to a significantly lower degree compared to the wild-type (31). In addition, cytotoxicity against HeLa cells was observed by infection with a BspR (a negative regulator for type III secreted proteins)-deficient *B. pertussis* strain (15). These results suggest that the T3SS has a significant role in the virulence of *B. pertussis*.

The production and the relative amounts of *bopB* mRNA of Tohama I cultured in SS AsA^−^ medium were significantly increased compared to those of Tohama cultured in SS medium, but this was not the case for BteA mRNA (Figs. 2, 5). It was reported that *bteA* and *bopB* genes are upregulated by BtrS (a sigma factor for type III secreted proteins) and downregulated by BspR (13, 15). It has also been reported that the amount of *bteA* mRNA was increased 2-fold in a BspR-deficient strain compared to the wild-type, whereas the amount of *bopB* mRNA was increased by 18-fold compared to the wild-type by RNA-seq (15). The *bopB* gene is localized in the T3SS apparatus locus (*bsc* locus) adjacent to the *btrS* gene. Since the *bteA* gene is separated from the *bsc* locus by >2.5 Mb (9), BtrS would preferentially regulate transcription of the *bopB* gene over that of the *bteA* gene.

The cytotoxicity against L2 cells was observed by the infection of Tohama I cultured in SS_1/20AsA _1/2CaA medium (Fig. 6). Before infection, we exchanged Ham’s F-12K medium for the SS_1/20AsA _1/2CaA medium. When L2 cells were infected with Tohama I in Ham’s F-12K medium, the cytotoxicity against L2 cells was not observed (Fig. S1). In order to establish oxidizing conditions in the tissue culture media, we used several oxidizing agents, i.e., hydrogen peroxide (H_2_O_2_), potassium ferricyanide(III) (K_3_[Fe(CN)_6_]), methyl 3-nitro-2-pyridinesulfenate (Npys-OMe), and 5,5’-dithiobis(2-nitrobenzoic acid) (DTNB). We also used another tissue culture medium, i.e., Minimum Essential Medium Eagle (MEM), because MEM contains no reducing agents, while F-12K, which is the medium for the L2 cells culture, contains a reducing agent, i.e., cysteine. Even when L2 cells were infected in F-12K medium containing the oxidizing agent or MEM, *B. pertussis* showed no cytotoxicity (Fig. S1). We therefore speculate that a substance which inhibits the secretion of type III secreted proteins from Tohama I was present in the Ham’s F-12K medium, or a substance that is necessary for the secretion of type III secreted proteins from Tohama I was absent in the Ham’s F-12K medium. We found that infection of the human alveolar epithelium cell line A549 with wild-type *B. pertussis* cultured in SS_1/20AsA _1/2CaA medium did not result in cytotoxicity (data not shown). For the observation of cytotoxicity against A549 cells, it is necessary to optimize the conditions for the production and secretion of type III secreted proteins in *B. pertussis*.

A previous study showed that cytotoxicity against the cell line J774A.1 was achieved by infection with wild-type *B. pertussis* 18323, but not by infection with a CyaA-deficient strain (34). This finding suggested that CyaA has the potential to provoke the cytotoxicity. However, we confirmed herein that the production and secretion of CyaA in a BteA-deficient strain (MGKU3) were not significantly different from those of the wild-type (data not shown). In our results, the amounts of LDH released from L2 cells infected with MGKU3 were greater than those of the T3SS-inactive strain (MGKU2), and no LDH release was detected from L2 cells infected with MGKU2 (Fig. 8). These results raised the possibility that an unknown type III effector other than BteA is involved in the cytotoxicity of *B. pertussis*.

Although we observed no significant difference in the growth rate of Tohama I between SS medium and SS_AsA^−^ medium (data not shown), the production and secretion of type III secreted proteins of Tohama I cultured in the SS_AsA^−^ medium were increased compared to those in the SS medium (Fig. 2). It had not been determined whether the expression of type III secreted proteins is regulated by reducing agents at the molecular level, and in this study, the production and secretion of CyaA and FHA in Tohama I cultured in SS medium were not significantly different from those in SS_AsA^−^ medium (Fig. 2). It was reported that there is no difference in Prn production by Tohama I cultured in modified SS medium compared to culture in modified SS medium containing dithiothreitol (DTT), a reducing agent (35). These results suggest that BvgAS-regulated virulence factors other than type III secreted proteins are not affected by reducing agents.

Ascorbic acid, urea, and glutathione (GSH) function as reducing agents in the epithelial lining fluid (ELF) of the human airway. The concentrations of ascorbic acid and urea in ELF in the upper respiratory tract are not significantly different from those in the lower respiratory tract. However, the concentration of GSH in the ELF in the upper respiratory tract is 100-fold lower than that in the lower respiratory tract. One of the possibilities is that GSH forms GS-SG (an oxidized from of GSH) in the ELF in the upper respiratory tract (36). It may thus be considered that the upper respiratory tract is more oxidized than the lower respiratory tract. It was reported that the transcription of genes encoding type III secreted proteins of *B. pertussis* recovered from mouse nasal lavage was higher than that from bronchoalveolar lavage (37). We thus speculate that the T3SS is important for *B. pertussis* to colonize the upper respiratory tract of the host.

In this study, the production and secretion of BopB by Tohama I cultured in the SS_AsA^−^ medium were significantly increased compared to those in SS medium (Fig. 2). Bacteria use redox-sensing proteins to resist oxidative stress (38). For example, OxyR (a member of the LysR family of transcriptional regulators) of *Escherichia coli* is converted to an active mode by the oxidation of intramolecular cysteine residues, which then increases the expression of antioxidants genes, e.g., hydroperoxidase I [*katG*, (39)]. Vfr (a member of the CRP/FNR superfamily) of *Pseudomonas* forms an active mode by retaining intramolecular cysteine residues as reducing states, and then increases the expression of type III secreted proteins (40). Btr is a regulator protein produced by *B. pertussis* and is homologous to Vfr (41). A cysteine cluster is located in the N-terminal domain of Btr, and a helix-turn-helix DNA binding domain is located in the C-terminal domain of Btr. Therefore, when *B. pertussis* is cultured in reducing agent-deprived SS medium, Btr might be oxidized and change its conformation, thereby increasing the expression of type III secreted proteins. Although the signal pathway that *B. pertussis* uses to increase the production of type III secreted proteins is unclear, it is possible that Btr regulates gene expressions for type III secreted proteins.

## ACKNOWLEDGMENTS

This work was supported in part by grants from the Ministry of Education, Culture, Sports, Sciences and Technology and the Japan Society for the Promotion of Science (KAKENHI), nos. 19K07542 (to A.A.), 17K08838 (A.K.), and 19K07561 (T.H.), and by a research grant from the Takeda Science Foundation in 2016 (A.K.). The funders had no role in the study design, data collection or analysis, the decision to publish, or the preparation of the manuscript.

---

**Fig. S1.**
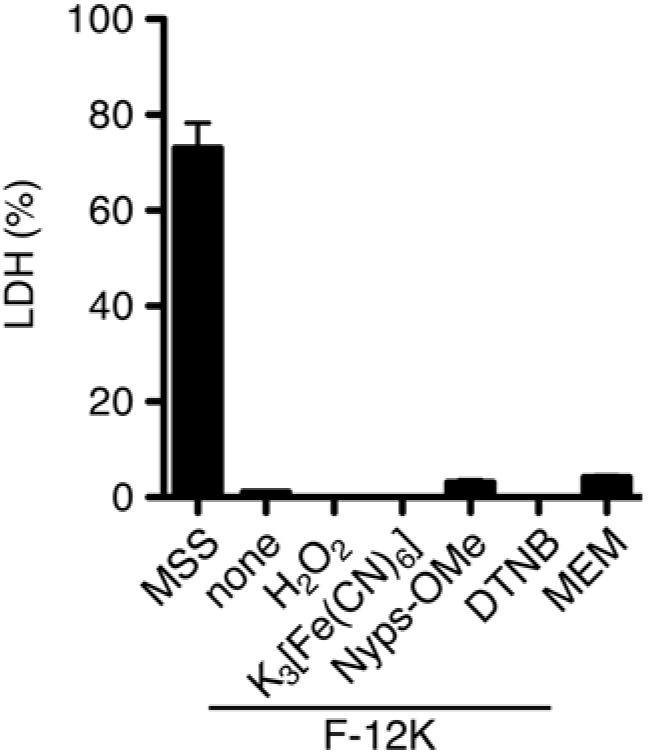
Cytotoxic activity of *B. pertussis* Tohama I in F-12K medium containing oxidizing agents. L2 cells were infected with *B. pertussis* Tohama I in the modified SS medium (SS_1/20AsA_1/2CaA, MSS). The amounts of ascorbic acid and casamino acids in MSS are one-twentieth of, and one-half of those in SS medium, respectively. L2 cells were also infected in F-12K medium (none), F-12K medium containing hydrogen peroxide (H_2_O_2_), potassium ferricyanide(lll) (K_3_[Fe(CN)_6_]), methyl 3-nitro-2-pyridinesulfenate (Npys-OMe) or 5,5’-dithiobis(2-nitrobenzoic acid) (DTNB). The final concentrations of each oxidizing agent were 1 mM, 1 mM, 10 *μM*, or 2 mM, respectively. Minimum Essential Medium Eagle (MEM) was also used. The cells were infected at an MOI of 500 for 3 hr. The amounts of LDH released into the extracellular medium from infected cells are shown, and the relative cytotoxicity (%) was determined as described in the Materials and Methods section. Error bars are the SEM from triplicate experiments. Experiments were performed at least three times, and representative data are shown.

